# The DNA methyltransferase 1 (DNMT1) acts on neurodegeneration by modulating proteostasis-relevant intracellular processes

**DOI:** 10.1101/2020.07.10.197442

**Authors:** Cathrin Bayer, Georg Pitschelatow, Nina Hannemann, Jenice Linde, Julia Reichard, Daniel Pensold, Geraldine Zimmer-Bensch

## Abstract

The limited regenerative capacity of neuronal cells requires tight orchestration of cell death and survival regulation in the context of longevity, as well as age-associated and neurodegenerative diseases. Subordinate to genetic networks, epigenetic mechanisms, like DNA methylation and histone modifications, are involved in the regulation of neuronal functionality, and emerge as key contributors to the pathophysiology of neurodegenerative diseases. DNA methylation, a dynamic and reversible process, is executed by DNA methyltransferases (DNMTs). DNMT1 was previously shown to regulate neuronal survival in the aged brain, whereby a DNMT1-dependent modulation of processes relevant for protein degradation was proposed as underlying mechanism. Functional proteostasis networks are a mandatory prerequisite for the functionality and long-term survival of neurons. Malfunctioning proteostasis is found, inter alia, in neurodegenerative contexts. Here, we investigated whether DNMT1 affects critical aspects of the proteostasis network by a combination of expression studies, life cell imaging and biochemical analyses. We found that DNMT1 negatively impacts retrograde trafficking and autophagy, both being involved in the clearance of aggregation-prone proteins by the aggresome-autophagy pathway. In line with this, we found that the transport of GFP-labeled mutant HTT to perinuclear regions, proposed to by cytoprotective, also depends on DNMT1. Depletion of *Dnmt1* accelerated HTT perinuclear HTT aggregation and improved the survival of cells transfected with mutant HTT. This suggests that mutant HTT-induced cytotoxicity is at least in part mediated by DNMT1-dependent modulation of degradative pathways.

## 1. Introduction

The limited regenerative capacity of neurons requires tight regulation of neuronal functionality and survival [1]. Epigenetic mechanisms of transcriptional control, like DNA methylation, catalyzed by DNA methyltransferases (DNMTs), as well as histone modifications, contribute to cell death regulation during development, aging and disease [2, 3]. DNMT1 is one of the main DNMTs expressed in the developing and adult brain. In addition to its function in dividing progenitors, DNMT1 catalyzes promoter as well as gene body cytosine methylation in post-mitotic and mature neurons [4-7]. Functionally, DNMT1 was already reported to promote the survival of a variety of maturing neurons, including excitatory neocortical and hippocampal neurons [8], photoreceptor cells and other types of neurons in the postnatal retina [9]. Besides, we found that DNMT1 is crucial for the survival of developing GABAergic inhibitory interneurons [10].

In contrast to its survival promoting function in developing neurons, we revealed a negative effect of DNMT1 on cortical interneuron longevity in the aged brain [11]. However, no cell death or survival-related genes were revealed to be targeted by DNMT1 function as an underlying cause. In turn, a significant enrichment of genes associated to proteostasis pathways were found to be suppressed by DNMT1 [10-12].

The proteostasis network ultimately impacts on long-term health of neurons. Its decline causing ineffective protein degradation can lead to neuronal death, and emerges to be implicated in age- and disease-related neurodegeneration [13]. Lysosomal degradation is of great importance for removing defective proteins or protein aggregates delivered by autophagy- or endocytosis-triggered endosomal pathways [14-16]. Hence, it is not surprising that lysosomal dysfunction is associated with neurodegenerative pathologies [17, 18]. Similarly, autophagy, as the major intracellular machinery for degrading aggregated proteins and damaged organelles, is implicated in the disease pathology of many neurodegenerative disorders [19, 20].

Numerous neurodegenerative diseases are characterized by aggregation prone proteins, which are suggested to cause or to contribute to the disease pathophysiology. For example, Huntington’s disease (HD) is caused by the expansion of CAG repeats coding for glutamine (Q) in exon 1 of the huntingtin gene. These polyQ repeats lead to misfolding of the mutant huntingtin (HTT) protein, which are highly prone to aggregate [21-25].

It is proposed that misfolded proteins are actively transported to a cytoplasmic juxtanuclear structure, called an “aggresome”, when the chaperone refolding system and the ubiquitin-proteasome degradation pathway are overwhelmed by the production of such misfolded proteins [26]. Aggresome formation is considered as a cytoprotective response serving to sequester potentially toxic misfolded proteins into pericentriolar inclusion bodies and facilitate their clearance by autophagy [27-34]. The aggresome-autophagy pathway, proposed to concertedly eliminate such aggregation-prone proteins like mutant HTT [28], is coupled to the retrograde microtubule-based transport to recruit both the aggregated proteins as well as molecular determinants of autophagic vacuole formation and lysosomes to pericentriolar cytoplasmic inclusion bodies [32-34]. In support of this, microtubule-disrupting agents prevent aggresome formation, and result in elevated polyglutamine toxicity [30, 35]. In the context of HD, it was shown that DNMTs mediate the mutant HTT-induced cytotoxic effects in cortical neurons, whereas the underlying mechanism is unknown. We have evidence that DNMT1 represses the expression of genes related to autophagy, perinuclear region and intracellular microtubule trafficking events, like endo-lysosomal transport, that is crucial for degradative proteostatis networks [12]. Indeed, we found accelerated retrograde endosomal transport and trafficking of endocytosed cargo fated for lysosomal degradation upon *Dnmt1* depletion [11]. As retrograde trafficking is central to the aggresome-autophagy pathway, and as neurons expressing polyglutamine-expanded *Htt* show improved survival when they form aggresomes [36], we here aimed to investigate whether DNMT1 is implicated in mutant HTT-induced cytotoxicity by acting on retrograde transport, aggresome formation and autophagy.

## 2. Results

### DNMT1 modulates retrograde trafficking

In previous studies, we have identified that DNMT1 transcriptionally controls a variety of endosomal, lysosomal and retrograde trafficking-associated genes in their expression, with most of them being repressed by DNMT1 in cortical neurons ([11, 12]; **Figure 1a, Supplementary Fig. S1a**). Functionally, we confirmed that DNMT1 slows down the velocity of retrograde transportation of endolysosomal compartments in cerebellar granule (CB) cells [11], which we frequently use as neuronal cell culture model [11, 12]. Here we verified, whether retrograde transport is likewise modulated by DNMT1 in neuroblastoma (N2a) cells, to assess whether the regulation of retrograde transport is a general, neuronal subtype-independent function of DNMT1. To this end, we transfected N2a cells with *Cd63-GFP* and *Lamp1-mCherry*-encoding plasmids, to label endosomes and lysosomes [37], both being involved in autophagy and endocytosis-dependent degradation [38]. We then analyzed the velocity of anterogradely and retrogradely moving CD63- and LAMP1-containing compartments upon *Dnmt1* siRNA knockdown by using life cell imaging (**Fig. 1 b-f**). Compared to control siRNA treatment, *Dnmt1* depletion resulted in significantly elevated velocities of retrogradely moving CD63- and LAMP1-positive particles, while the anterograde transportation was not affected (**Fig. 1 d, f**). In line with this, the overall distances of retrogradely but not anterogradely transported CD63- and LAMP1-positive compartments were found significantly increased after Dnmt1 knockdown (**Supplementary Fig. S1 b, c**). Together, this suggests that DNMT1 selectively influences retrograde trafficking across different neuronal model systems.

**Figure 1.**
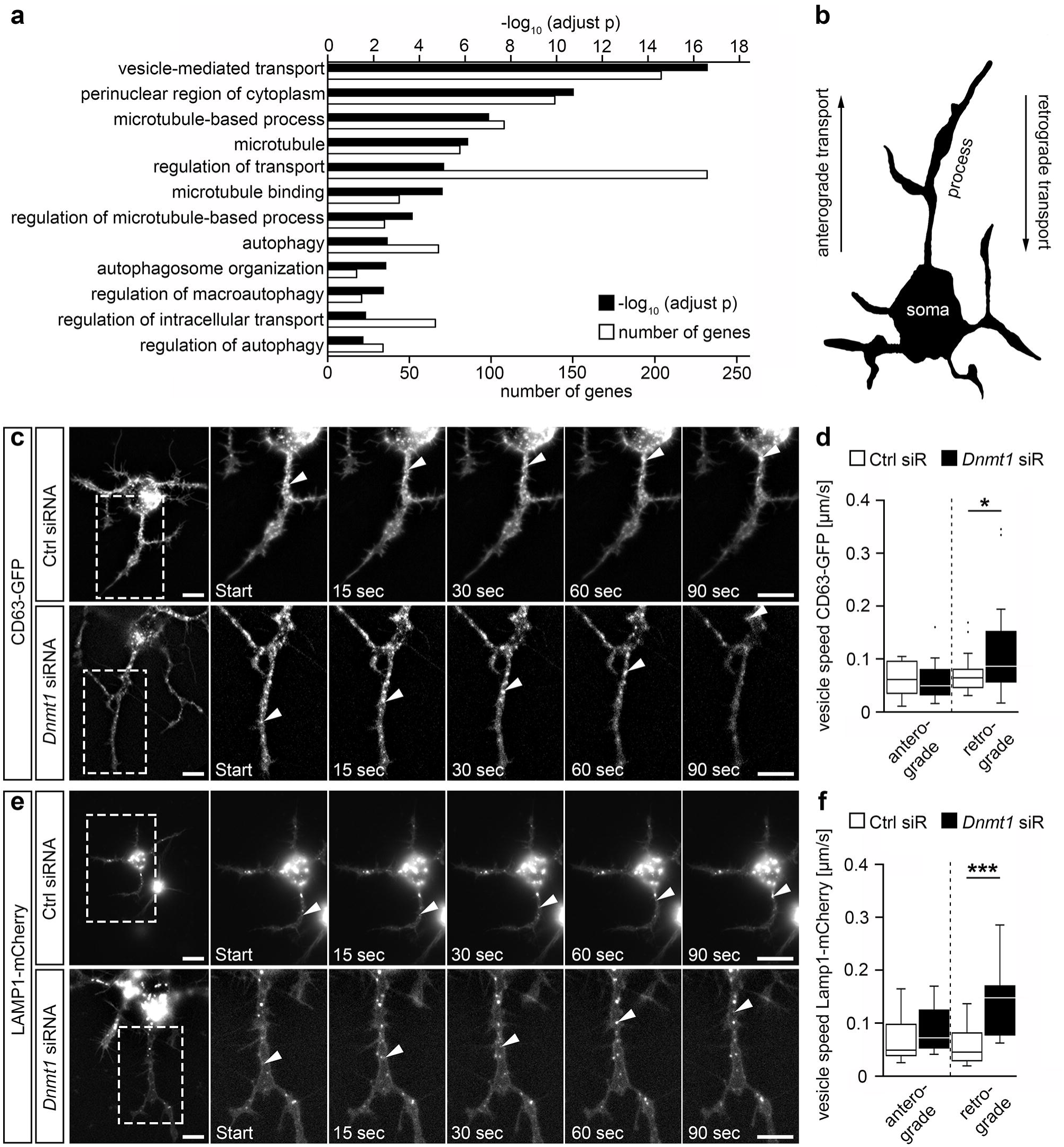
DNMT1 acts on dynamics of retrograde vesicle transport. (**a**) GO terms found significantly enriched for the genes, that were up-regulated upon *Dnmt1* deletion in adult *Pvalb-Cre* expressing cortical interneurons determined by RNAsequencing after FAC-sorting of *Pvalb-Cre*/*tdTomato*/*Dnmt1* WT and KO mice (*P* < 0.05, Benjamini-adjusted, n = 9 WT and n = 12 KO mice). (**b**) Schematic representation of a N2a cell with implied axonal transport directions. **(c)** Representative microphotographs of the tracking of CD63-GFP-labeled vesicles during live cell imaging in N2a cells transfected with *CD63-GFP* plasmid and Ctrl siRNA (n = 14 cells and 34 vesicles) or *Dnmt1* siRNA (n = 14 cells and 38 vesicles). The vesicle velocity is quantified in (**d**) (Two-sided Student’s t-test, * P < 0.05). (**e**) Representative microphotographs of the tracking of LAMP1-mCherry-positive vesicles during live cell imaging in N2a cells transfected with *Lamp1-mCherry* plasmid and Ctrl siRNA (n = 20 cells and 38 vesicles) or *Dnmt1* siRNA (n = 17 cells and 37 vesicles). The vesicle velocity is quantified in (**f**) (Two-sided Student’s t-test, *** P < 0.001). Ctrl = control, KO = knockout, LAMP1 = Lysosomal-associated membrane protein 1, N2a = neuroblastoma cells, siR = siRNA, WT = wildtype. Scale bars: 10 μm in (c, e).

### DNMT1-mediated retrograde trafficking to perinuclear regions influences mutant HTT-induced cytotoxicity

Degradative proteostasis networks rely on retrograde trafficking in many regards. Apart from trafficking of endosomes to degradative lysosomes or secretory compartments [39], retrograde transport is crucial for autophagic clearance [40]. Indeed, defects in the retrograde transport of autophagosomes are involved in disease-related neurodegeneration [40, 41]. Moreover, the aggresome-autophagy pathway relies on microtubule-based retrograde transportation [26]. When the ubiquitin proteasome system is overwhelmed with aggregation-prone proteins, the aggresome-autophagy pathway sequesters misfolded proteins into pericentriolar inclusion bodies called “aggresomes” and facilitates their clearance [33, 34]. Aggresome formation is suggested to improve neuronal survival upon expression of polyglutamine-repeat containing proteins like mutant HTT, that escape proteasome-dependent degradation [42].

In line with this, we observed numerous N2a and CB cells forming GFP aggregates in the perinuclear region upon transfection with a mutant *Htt-GFP* containing plasmid and inhibition of proteasomal degradation with MG-132. The utilized plasmid encodes for mutant HTT-GFP, leading to the expression of the GFP-labeled exon 1 of the mutated Huntingtin protein with a total of 103 glutamine repeats. While HTT-GFP was initially distributed equally in the cytosol at the beginning of the life cell imaging, a localized and intense GFP spot can be seen in numerous cells already 6 h of life cell imaging (**Fig. 2 a, b; Supplementary Fig. S2 a, c**), indicative of aggresome formation. The perinuclear localization of the HTT-GFP aggregate can be seen clearly for CB cells (**Fig. 2d, Supplementary Fig. S2 c**). In support of a survival promoting function of aggresome formation, we found increased survival rates for N2a cells, that formed visible HTT-GFP aggregates, compared to the cells with no apparent aggregation of the mutant HTT-GFP protein (**Supplementary Fig. S2a, b**). Similar observations in matters of improved survival upon aggregate formation were made for CB cells (**Supplementary Fig. S2c, d**).

**Figure 2.**
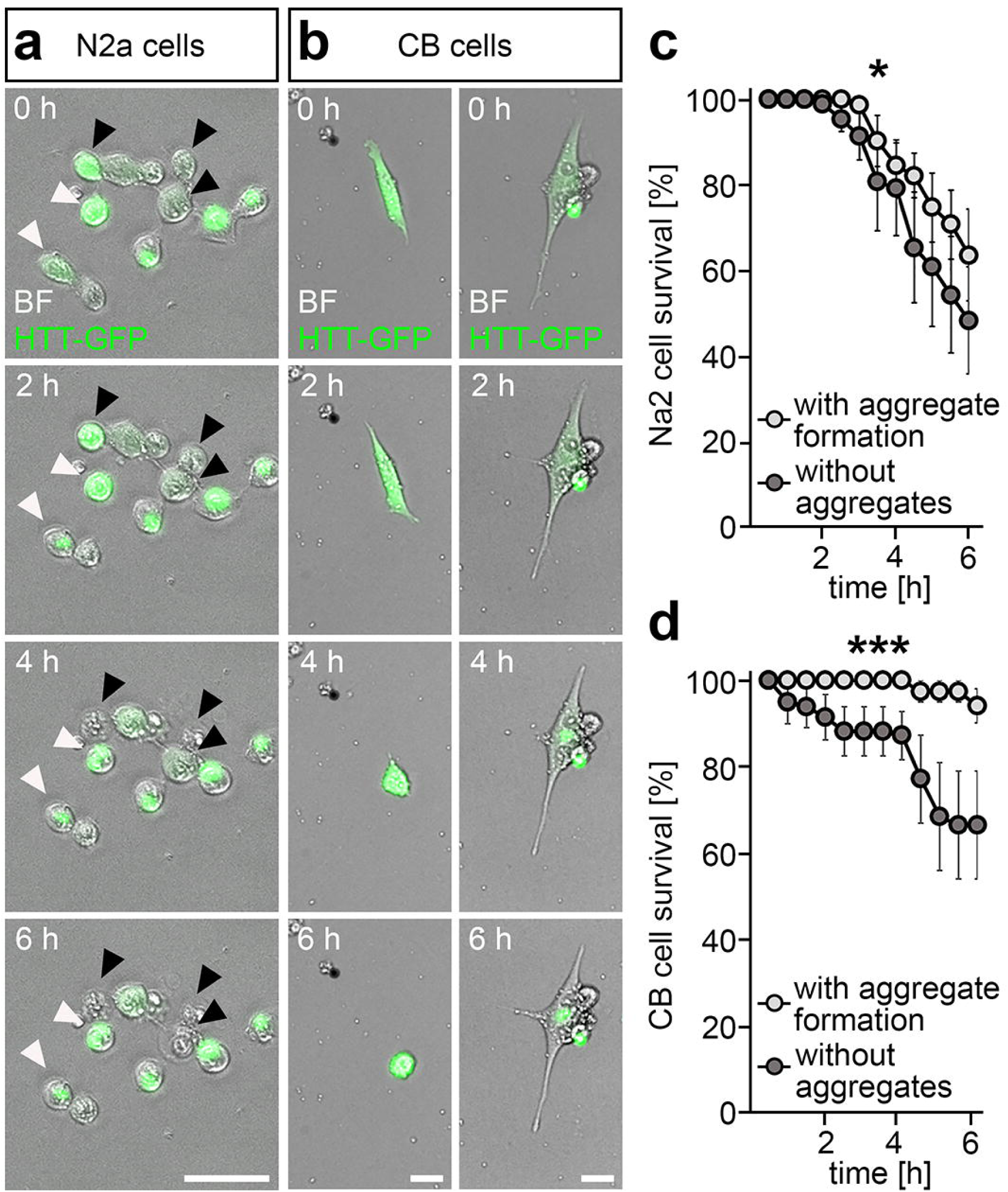
Aggregate formation of mutant HTT is linked to a higher rate of cell survivability. (**a**) Representative microphotographs of N2a cells expressing GFP-labeled mutant HTT (green) showing higher survivability in cells with aggregate formation (white arrows) than in cells without aggregates (black arrows). (**b**) Representative microphotographs of CB cells expressing GFP-labeled mutant HTT (green) showing higher survivability in a cell with aggregate formation (right column) than in a cell without aggregates (left column). (**c**) Survival rates of N2a cells expressing GFP-labeled mutant HTT over the time course of 6 h in the HTT-cytotoxicity assay discriminating between cells with aggregate formation (N = 4 experiments, n = 113 cells) and cells without aggregates (N = 4 experiments, n = 64 cells; Two-way ANOVA, * P < 0.05). (**d**) Survival rates of CB cells expressing GFP-labeled mutant HTT over the time course of 6 h in the HTT-cytotoxicity assay discriminating between cells with aggregate formation (n = 27 cells) and cells without aggregates (n = 44 cells; Two-way ANOVA, * P < 0.05). BF = brightfield, CB = cerebellar granule cells, HTT = Huntingtin, N2a = neuroblastoma cells. Scale bars: 50 μm in (a), 20 μm in (c).

Interestingly, DNMTs were reported to be involved in mediating the cytotoxic effects of mutant HTT, whereas the underlying mechanism is unknown. As we found that DNMT1 restricts effective intracellular retrograde trafficking and represses genes relevant for the formation of the perinuclear region and microtubule-based processes (**Supplementary Fig. S1a**), we asked whether DNMT1 could interfere with HTT-GFP aggregation in the perinuclear region and through this, promote mutant HTT induced cytotoxicity.

To this end, we used life cell imaging to monitor the survival of mutant HTT-GFP expressing N2a and CB cells, in which proteasomal degradation was blocked after transfection with *Dnmt1* or control siRNA (**Fig. 3**). While mutant HTT protein expression resulted in a severe reduction of cell survival over a time period of 12 hours for both N2a and CB cells, the mutant HTT-induced cytotoxicity was significantly decreased in both cell lines upon knockdown of *Dnmt1*. *Dnmt1* depletion caused improved survival rates, similar to what was reported by Pan et al. ([43], **Fig. 3**).

**Figure 3.**
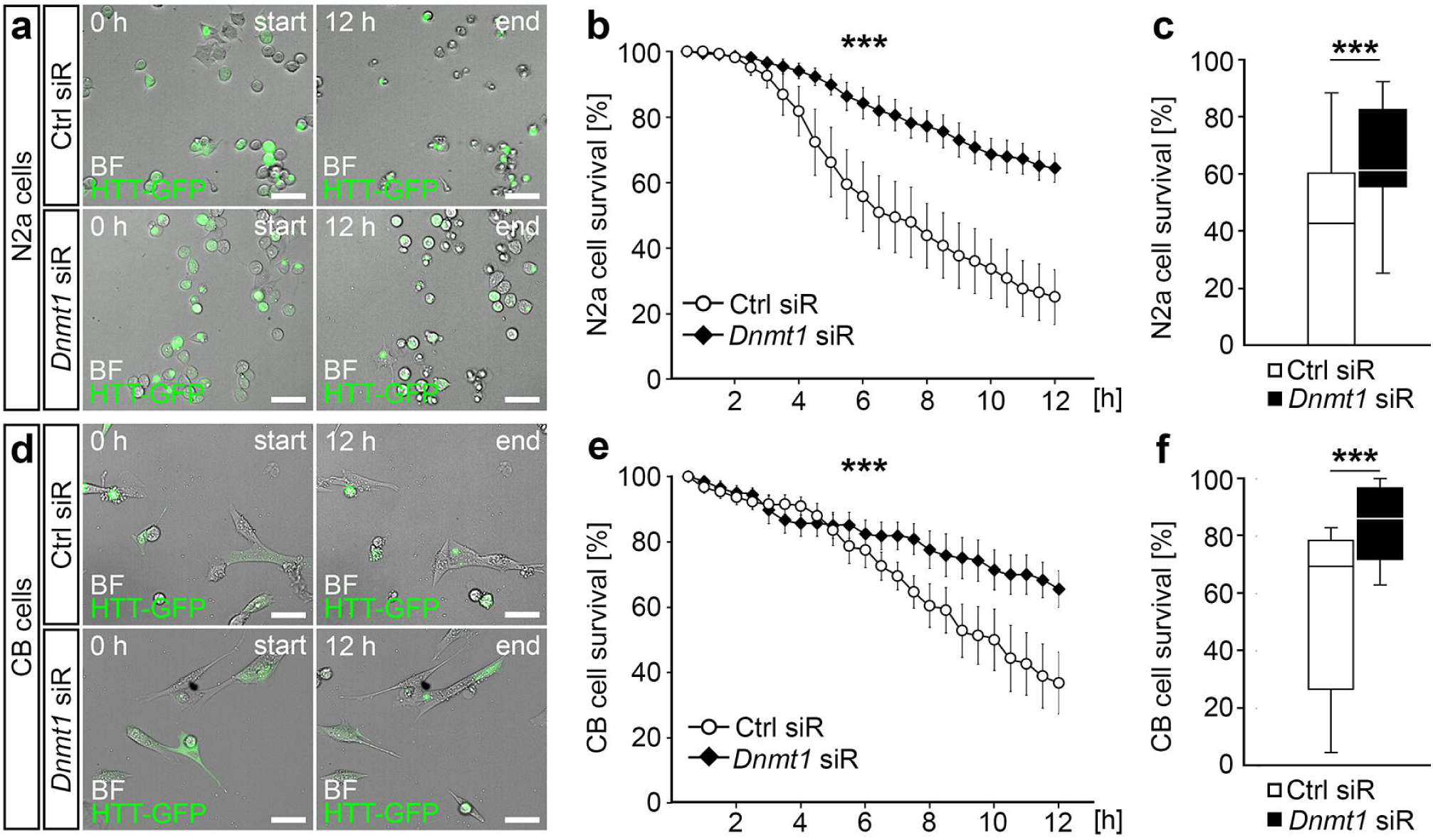
*Dnmt1*-knockdown ameliorates the neuronal cell survival upon mutant HTT-induced cytotoxicity. (**a**) Representative microphotographs of start and end points of 12 h in the HTT-cytotoxicity assay in N2a cells expressing GFP-labeled mutant HTT (green). Cells were transfected with either Ctrl or *Dnmt1* siRNA. (**b**) Cell survival rates of N2a cells expressing GFP-labeled mutant HTT and transfected with Ctrl siRNA (N = 4 experiments, n = 274 cells) or *Dnmt1* siRNA (N = 4 experiments, n = 378 cells) over the time course of 12 h in the HTT-cytotoxicity assay (Two-way ANOVA, *** P < 0.001). (**c**) Comparison of the survival rates of cells analyzed in (**b**) at the 12 h timepoint (Two-sided Student’s t-test, *** P < 0.001). (**d**) Representative microphotographs of start and end points of 12 h in the HTT-cytotoxicity assay in CB cells expressing GFP-labelled mutant HTT (green). Cells were transfected with either Ctrl or *Dnmt1* siRNA. (**e**) Cell survival rates of CB cells expressing GFP-labeled mutant HTT and transfected with Ctrl siRNA (n = 100 cells) or *Dnmt1* siRNA (n = 132 cells) over the time course of 12 h in the HTT-cytotoxicity assay (Two-way ANOVA, *** P < 0.001). (**f**) Comparison of the survival rates of cells analyzed in (**e**) at the 12 h timepoint (Two-sided Student’s t-test, *** P < 0.001).N2a = neuroblastoma cells, CB = cerebellar granule cells, Ctrl = Control, HTT = Huntingtin, siR = siRNA, BF=brightfield. Scale bars: 50 μm in (a, d).

Next, we aimed to know whether the implication of DNMT1 in mediating the mutant HTT-induced cytotoxicity depends on the DNMT1-dependent modulation of retrograde trafficking. To this end, we aimed to measure the velocity of HTT-GFP aggregate formation. For this, we focused on CB cells. Due to their larger soma size, the dynamics and the temporal course of perinuclear HTT aggregation can be clearly identified by life cell imaging, in contrast to aggregation in the smaller-sized and roundish N2a cells. In line with our findings that *Dnmt1* siRNA application increased the velocity of retrograde transport [11], we found that the formation of HTT-GFP aggregates was significantly faster upon siRNA mediated depletion of *Dnmt1* in CB cells (**Fig. 4**). This indicates that DNMT1 is involved in mutant HTT-induced cytotoxicity via modulation of aggresome formation by acting on retrograde transportation.

**Figure 4.**
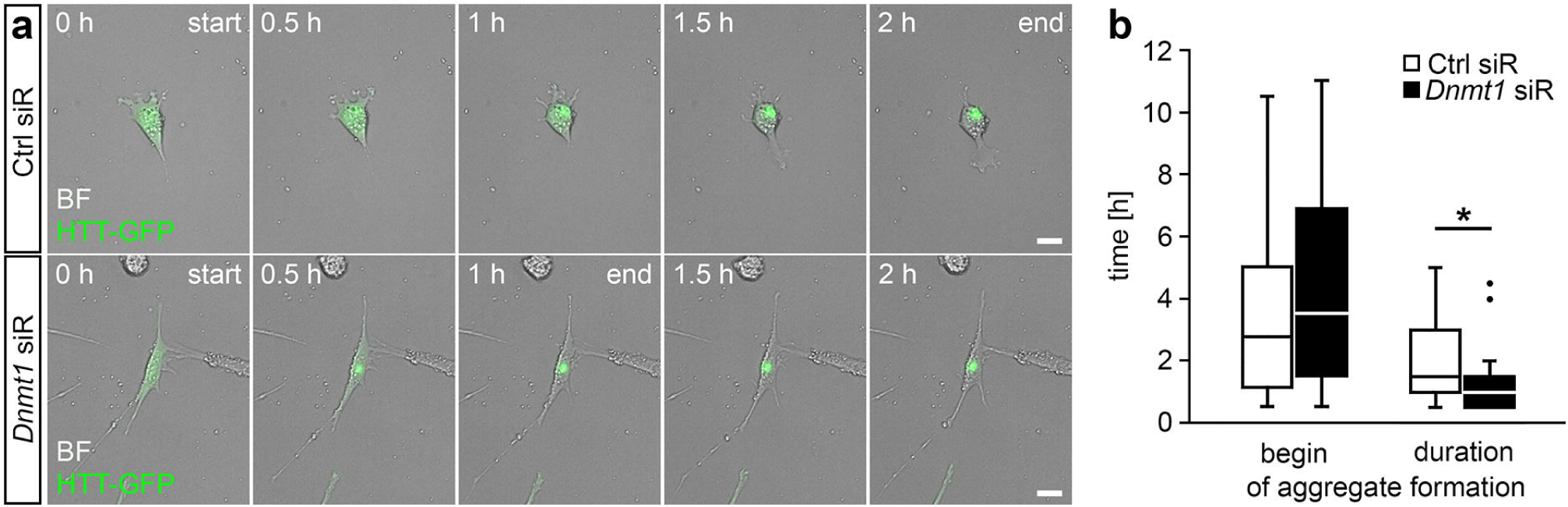
*Dnmt1*-knockdown leads to a faster aggregate formation of mutant HTT in CB cells. (**a**) Representative microphotographs of CB cells expressing GFP-labeled mutant HTT (green) and transfected with either Ctrl or *Dnmt1* siRNA, showing the temporal differences in the aggregate formation of mutant HTT. (**b**) Quantification of the begin and duration of aggregate formation depicted in (**a**) for cells transfected with Ctrl siRNA (n = 27 cells) or *Dnmt1* siRNA (n = 33 cells; Two-sided Student’s t-test, * P < 0.05). BF = bright field, Ctrl = control, HTT = Huntingtin, siR = siRNA. Scale bars: 20 μm in (a).

### DNMT1 acts on autophagy

The accumulation of aggregated proteins is suggested to induce an autophagic response to concertedly eliminate aggregation-prone proteins. To this end, molecular determinants of autophagic vacuole formation and lysosomes have been reported to be recruited to the pericentriolar cytoplasmic inclusion bodies, which relies on microtubule-dependent retrograde trafficking and which we found to be decelerated by DNMT1 [28].

Interestingly, we found autophagy-related genes collected in Figure 5a to be repressed by DNMT1 as well. Hence, we next examined whether DNMT1 function modulates autophagy. Autophagy starts with the formation of a phagophore, a double membrane that encloses and isolates the cytoplasmic components, which leads to the generation of autophagosomes. Autophagosomes fuse with multivesicular bodies (MVB)/late endosomes, which matured from early endosomes, generating an amphisome. The amphisomes then get delivered to lysosomes, fusing to autolysosomes for degradation of the cargo ([38]; **Fig. 5 b**). The formation of autophagic vacuoles or autophagosomes is accompanied by the conjugation of the cytosolic microtubule-associated protein 1 light chain 3B-I (LC3B-I) with phosphatidylethanolamine, which results in the formation of the membrane-associated LC3B-II [44]. We monitored alterations in LC3B-II levels using Western blot in N2a cells, that were either treated with control or *Dnmt1* siRNA (**Fig. 5 c**). Autophagy was triggered in these cells by incubation of neurons in starving medium, resulting in elevated LC3B-II levels upon *Dnmt1* siRNA treatment (**Fig. 5 c, d**). Additionally, we added bafilomycin A1, which causes the inhibition of the fusion of autophagosomes and lysosomes. This leads to an impaired degradation and an accumulation of LC3B-II positive autophagosomes, which were likewise increased after *Dnmt1* siRNA treatment (**Fig. 5 c, d**). DNMT1 protein levels were analyzed as well, showing the clear reduction of the protein upon treatment with the respective siRNA (**Fig. 5 c, d**) and confirming the siRNA mediated knockdown.

**Figure 5.**
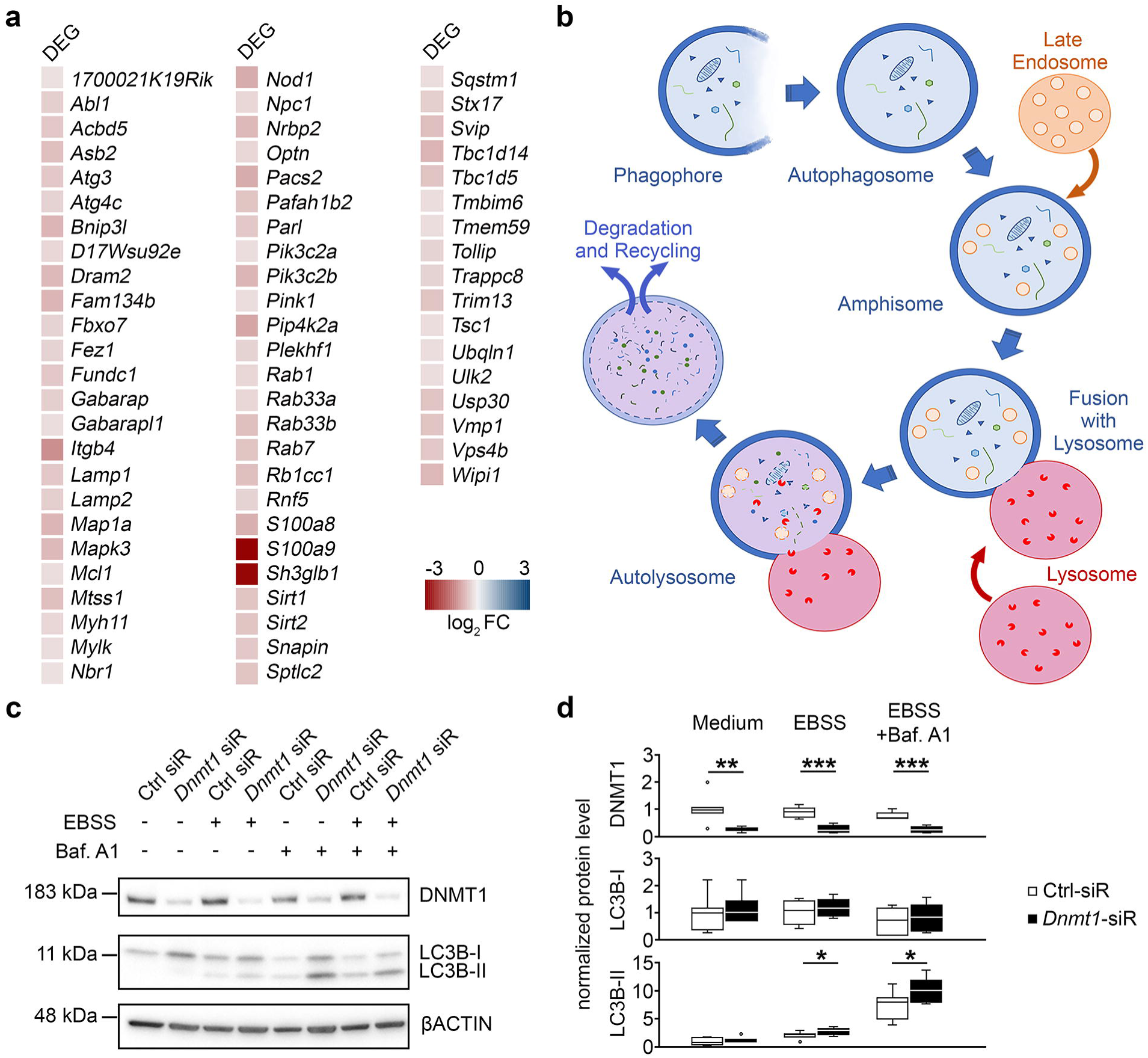
DNMT1 has an impact on autophagy. (**a**) Heat-map of significantly up-regulated genes associated to the GO term *autophagy* in FAC-sorted cortical interneurons of *Pvalb*-*Cre*/*tdTomato*/*Dnmt1* KO mice compared to cells of *Pvalb*-*Cre*/*tdTomato*/*Dnmt1* WT mice. Expression level determined by RNA-sequencing (*P* < 0.05, Benjamini-adjusted, n = 9 WT and n = 12 KO mice). (**b**) Schematic illustration of the autophagy pathway beginning with the formation of an autophagosome and depicting the transition to an amphisome, which fuses with a lysosome to form the autolysosome, where particles are degraded. (**c**) Protein levels of DNMT1, LC3B-I and -II in Ctrl and *Dnmt1* siRNA treated N2a cells in the autophagy assay analyzed by Western blot. Cells were cultured in culture medium or starving solution (EBSS) and partly treated with an autophagy inhibitor (Bafilomycin A1). (**d**) Quantification of protein levels of the conditions shown in (**c**) normalized to bACTIN and in relation to the control (N = 7 experiments; Two-sided Student’s t-test, * P < 0.05, ** P < 0.01, *** P < 0.001). DEG = differentially expressed genes, FC = foldchange, bACTIN = beta-actin, Baf. A1 = bafilomycin A1, Ctrl = control, EBSS = Earle’s buffered salt solution, LC3B-I/II = Microtubule-associated protein 1 light chain 3B-I/II, N2a = neuroblastoma cells, siR =siRNA.

In sum, DNMT1 seems to modulate autophagy. Collectively, these data suggest that DNMT1 is involved in the mutant HTT-induced cytotoxicity by acting on degradative pathways including retrograde transportation, aggresome formation and autophagy.

## 3. Discussion

Innumerable potential mechanisms are proposed to be involved in neurodegeneration, such as defects in protein homeostasis, impaired protein degradation, alterations in gene expression and transcriptional regulation, in addition to mitochondrial dysfunction [45]. However, a prerequisite to improve early diagnosis and to develop disease-modifying therapy strategies is the detailed understanding of the interplay and hierarchies of the pathophysiological mechanisms.

Epigenomic remodeling call increasing attention in the field of neurodegeneration, proposed to mediate the initiation and progression of neurodegenerative disorders, and to potentially serve as novel targets for therapeutic interventions [46]. Epigenetic transcriptional control is known to orchestrate diverse aspects of neuronal physiology in the developing and adult brain, being essentially implicated in the regulation of neuronal survival and function [2, 3, 47]. Epigenomic signatures are moreover dynamically reconfigured in the aging brain, and suggested to mediate age-related alterations, like the loss of synapses and even particular neuronal subpopulations [3, 11]. Hence, it is not surprising that epigenetic dysregulation has been proposed to be implicated in events underlying neuronal dysfunction and neuronal cell death in diverse neurodegenerative diseases including Huntington’s disease (HD) [47-49].

HD is caused by the expansion of CAG repeats coding for glutamine (Q) in exon 1 of the huntingtin gene [50, 51]. The mutated HTT protein has been reported to act on a wide range of epigenetic signatures, including histone modifications (i.e. acetylation, methylation and ubiquitination), and DNA methylation marks [49, 52]. These alterations can be associated to transcriptional dysregulation, accounting as a major pathogenic mechanism of HD and an early event in HD pathology, preceding the onset of neuronal cell death [52, 53]. In line with this, DNA methylation signatures were found to be changed in HD patients and HD animal models, correlating with alterations in gene expression (reviewed in [47]).

It was described by Pan et al., 2016 [43] that DNMTs, catalyzing DNA methylation, promote mutant HTT-induced cytotoxicity, whereas the underlying mechanism remains unknown. We found that DNMT1 is implicated in the age-related neurodegeneration, underlining the importance of DNMT function and DNA methylation for neuronal longevity regulation [11]. Thereby, DNMT1 does not seem to target cell death- or survival-associated genes to modulate neuronal survival. Instead, DNMT1 influences neuronal homeostasis by regulating the expression of proteostasis-related genes, including endocytosis, endosome, lysosome and autophagy-related genes [11, 12].

A decline in protein homeostasis is known to contribute to numerous neurodegenerative disorders [13]. Due to the limited regenerative capacity of neurons, clearance of defective proteins or protein aggregates by proteolytic degradation in lysosomes is of great importance for neuronal homoeostasis [54]. Lysosomal dysfunction as well as defective autophagy are implicated in neurodegenerative disorders [55-61].

In line with the transcriptional repression of autophagy-related genes, we here functionally verified that DNMT1 negatively regulates autophagy. Further, we found that DNMT1 acts as a brake on retrograde transportation of endo-lysosomal compartments. Apart from the transportation of endosomes to degradative lysosomes or secretory compartments [39], retrograde trafficking is crucial for autophagic clearance [40]. Defective retrograde trafficking of autophagosomes contributes to disease-related neurodegeneration [40, 41]. Besides, microtubule-based retrograde transportation is essential for the aggresome-autophagy pathway [26]. This pathway sequesters misfolded proteins into pericentriolar inclusion bodies called “aggresomes” and facilitates their clearance, suggested to compensate for defects in the ubiquitin proteasome-dependent degradation of aggregation-prone proteins [33, 34]. Aggresome formation is proposed to improve neuronal survival upon expression of polyglutamine-repeat containing proteins, like mutant HTT, that escape proteasome-dependent degradation [42]. It was shown that inclusion formation reduced the amount of mutant HTT in other areas of the cell, being associated with increased cell survival [36, 62]. This is in line with our findings, showing elevated survival rates of mutant Htt-expressing cells upon aggregate formation. Interfering with aggresome formation, which relies on microtubule-based retrograde trafficking, by inhibiting microtubule polymerization or impairing dynein motor function leads to decreased viability of cells expressing disease proteins [30]. Interestingly, we found that the retrograde transport and the aggregation of mutant HTT-GFP are accelerated and the HTT-induced cytotoxicity is reduced when *Dnmt1* is depleted. Together, this indicates that DNMT1 acts on aggresome formation by modulating retrograde transportation to perinuclear regions. In addition to the transport of aggregated proteins to perinuclear regions, retrograde trafficking is further suggested to recruit autophagy-relevant compartments like autophagosomes and lysosomes [63]. Indeed, there is accumulating evidence that aggresomes are substrates for autophagy [27-31], even being proposed to concentrate aggregated proteins for more efficient autophagic degradation [28, 64, 65]. As DNMT1 affects both retrograde transportation and autophagy, DNMT1 could indirectly promote mutant HTT-induced cytotoxicity by lowering the efficiency of the aggresome-autophagy pathway.

However, we cannot exclude that DNMT1 directly interacts with mutant and/or wild-type HTT and possibly mediates changes in epigenomic and transcriptional remodeling. Indeed, HTT is assumed to be a multi-functional protein, that participates in diverse cytosolic as well as nuclear processes [47]. Apart from its putative function in vesicular trafficking, HTT is localized in the nucleus and modulates transcription by binding to transcription factors. Moreover, HTT affects the function of epigenetic key players like RE1-silencing transcription factor (REST) and the multi-subunit polycomb repressive complex 2 (PRC2) (reviewed in [47]). PCR2 is an epigenetic silencer complex catalyzing repressive H3K27 trimethylations [66]. Rising numbers of polyglutamine-repeats in mutant HTT were shown to progressively increase the histone H3K27 tri-methylase activity of PRC2 *in vitro* [67]. This is in line with the finding of genome-wide changes in H3K27me3 signatures upon expression of mutant *Htt* in embryonic stem cells [68]. The connection between mutant HTT and PRC2 is further reinforced by changes in death-promoting proteins upon induced PRC2-deficiency in striatal medium spiny neurons (MSNs). Moreover, transcriptional changes of transcription factors, which are normally suppressed in MSNs, were observed, accompanied by progressive and fatal neurodegeneration [69]. In addition to that, loss of PRC2 in forebrain neurons elevates the expression of genes involved in HD [69].

Although direct interaction of HTT and DNMT1 were to our best knowledge not described so far, mutant HTT could as well indirectly interfere with DNMT1 function, as DNMT1 has been proposed to interact with PRC2 [70-73]. Moreover, certain histone modifications can prevent or promote DNA methylation [74], whereby DNMT1 function could be secondarily affected by mutant HTT-induced changes of the histone code. Non-coding RNAs (ncRNAs) likewise exert transcriptional control in part by recruiting or preventing the binding of proteins/complexes involved in setting up DNA methylation and histone modification marks [75]. Mutant HTT modulates REST function, which controls the expression of diverse long non-coding RNAs (lncRNAs) and micro RNAs (miRNAs), several of which are found dysregulated in HD [76]. Hence, DNMT1 function and DNA methylation targeting could be affected by the altered repertoire of ncRNAs. LncRNAs with critical functions in neuronal development [75] are known to regulate the binding of DNMTs to the DNA, thereby being involved in mediating site-specific methylation [77, 78].

Hence, to judge the potential of epigenetic mechanisms as targets for therapeutic interventions for neurodegenerative diseases like HD, we need to draw a conclusive picture. It is indispensable to shed light on the hierarchy of mutant HTT-induced epigenomic remodeling, and the function these epigenetic modifications have in healthy neurons. Moreover, we have to clearly dissect the primary effects from the compensatory responses of neurons when facing neurodegeneration. Here, we added up another part of the puzzle providing evidence that DNMT1-dependent transcriptional control lowers the efficiency of degradative processes, which could contribute to the polyQ-mediated cytotoxicity. In that regard, the decrease seen in *Dnmt1* expression in HD STHdhQ111 cells [79], and the diminished Dnmt1 expression observed in the striatum and cortex of N171-82Q transgenic HD mice may be interpreted as counterregulatory mechanisms of the cells to improve the efficacy of mutant HTT clearance.

## 4. Materials and Methods

### Cell lines and cell culture

Neuroblastoma (N2a) cells (ATCC: CCL-131) were cultured in Dulbecco’s modified Eagle’s medium with high glucose, GlutaMAX supplement and pyruvate (DMEM, #31966-021, Gibco), supplemented with 2% fetal bovine serum (FBS, Biowest), 100 U/mL penicillin (Gibco), 100 µg/mL streptomycin (Gibco) at 37°C, 5% CO2, and 95% relative humidity. The cells were splitted after reaching a confluence of 80-90%.

Cerebellar granule (CB) cells [80] were cultured in Dulbecco’s modified Eagle’s medium with high glucose (DMEM, #41965-039, Gibco) supplemented with 10% FBS (Biowest), 1% GlutaMAX, 24 mM KCl, 100 U/mL penicillin and 100 µg/mL streptomycin incubated at 33°C, 5% CO2 and 95% relative humidity. The cells were splitted after reaching a confluence of 75%.

### Transfection with siRNA and co-transfection with siRNA and plasmid DNA

Transfection of cells with siRNA was performed via lipofection using Lipofectamine 2000 or Lipofectamine 3000 (Thermo Fisher Scientific), according to the manufacturer’s protocol and as described in Zimmer et al. (2011) [81]. Mouse *Dnmt1* siRNA (30 nM; #sc-35203, Santa Cruz Biotechnology) or control siRNA (15 nM; Block-iT Alexa Fluor red (#14750100) or Block-iT green (#2013) fluorescent oligo, Invitrogen) were applied for 5 h in antibiotic- and serum-free Opti-MEM I Reduced Serum Medium (Thermo Fisher Scientific) and cells were then grown in respective cell culture media. For co-transfection with 30 nM *Dnmt1* siRNA (#sc-35203, Santa Cruz Biotechnology, USA), 15 nM control siRNA (Block-iT Alexa Fluor red (#14750100) or Block-iT green (#2013) fluorescent oligo, Invitrogen) and plasmid DNA (pLAMP1-mCherry, 200 ng/µL; CD63-pEGFP C2, 200 ng/µL; 1x GFP-pEGFP N3-HTT, 260 ng/µL), Lipofectamine 2000 (Thermo Fisher Scientific) was applied as described in the protocol of the manufacturer. Antibiotic- and serum-free Opti-MEM I Reduced Serum Medium (Thermo Fisher Scientific) were used for preparation of transfection reagents and dilutions. Co-transfections were performed for 24 h prior to life cell imaging.

### Monitoring of endo-lysosomal vesicle trafficking

To monitor the endo-lysosomal trafficking in N2a cells, 71 cells per mm^2^ were seeded in wells of a 24-well imaging plate (#0030741021, Eppendorf), that was previously coated with 19 µg/mL laminin (Sigma-Aldrich, Germany) and 10 µg/mL poly-L-lysine (Sigma-Aldrich) in Gey’s Balanced Salt Solution (GBSS). The coating solution was applied and incubated for 30 min at 37 °C. Subsequently, coverslips were washed once with sterile water and let dry prior to cell seeding. Cells were cultivated in 500 µl cell culture medium for 24 h using the above-mentioned culture conditions before co-transfection with siRNAs and plasmid DNA (pLAMP1-mCherry, CD63-pEGFP C2) was performed. Vesicle trafficking was captured via live cell imaging after 24 h of co-transfection.

### Huntingtin-cytotoxicity assay

For monitoring the mutant HTT-induced cytotoxicity, cells were seeded with densities of 55 cells per mm^2^ for CB cells and 137 cells per mm^2^ for N2a cells in a 24-well cell culture plate and were incubated for 24 h in 500 µl cell culture medium prior to co-transfection of siRNA and plasmid DNA (1x GFP-pEGFP N3-HTT). 21 h after transfection, cells were incubated with the proteasome blocker carbobenzoxy-L-leucyl-L-leucyl-L-leucinal (1 µM, MG-132, Sigma-Aldrich). After 2.5 h cells were additionally treated with the synthesis blocker cycloheximide (10 µg/mL, Sigma-Aldrich). Both inhibitors were diluted in phenol red-free cell culture medium (#21063-029, Gibco). Cells were then monitored using live cell imaging for either 6 and 12 h.

### Autophagy assay

To measure changes in autophagy, N2a cells were seeded at a density of 112 cells per mm^2^ in a 6-well cell culture plate in 2 mL culture medium, and were incubated for 24 h prior to siRNA transfection. Afterwards, cells were incubated for 2 h at 37 °C in 1 mL normal culture medium, supplemented with 200 nM Bafilomycin A1 (#sc-201550, Santa Cruz Biotechnology) or culture medium supplemented with DMSO. Subsequently, the medium was changed to either fresh culture medium with respective supplements (200 nM Bafilomycin A1 or DMSO) or to Earle’s balanced salt solution (EBSS, #14155-048, Gibco) supplemented with 200 mg/L MgSO4 and 200 mg/L CaCl2 with respective supplements (200 nM Bafilomycin A1 or DMSO). After 2 h the cells were washed and harvested in 1x PBS by rinsing the cells off the culture plate and centrifugation at 4°C for 3 min at 800 rcf. The supernatant was discarded and the cell pellet was lysed in RIPA lysis buffer (50 mM Tris-HCl (pH 8.0), 150 mM NaCl, 0.5% NP-40, 0.1% SDS, 0.1% sodium deoxycholate, freshly supplemented with 1 mM phenylmethylsulphonyl fluoride (PMSF, Serva) and 1 µg/mL Leupeptin (Serva)) for 10 min on ice. The lysate was treated twice for 30 sec respectively in an ultrasonic bath (100% power) and was later centrifuged at 4°C for 20 min at 21,000 rcf. The supernatant was transferred into a new tube and the pellet was discarded. Samples were prepared for Western blot as described below.

### Protein preparation, Western blotting and detection

The protein concentration of lysates was determined using the QuBit 4 fluorometer (Invitrogen). Samples with a protein amount of 30 µg were boiled in 4x sample buffer (200 mM Tris pH 6.8, 4% SDS, 10% β-mercaptoethanol, 40% glycerol and 0.002% bromophenol blue) at 95°C for 10 min, and subsequently stored at -20°C. The prepared protein samples were separated on a ready-to-use 4–12% gradient gel (Serva) using 1x SDS-running buffer (Laemmli buffer, Serva) following the manufacturer’s instructions. Proteins were transferred to a nitrocellulose membrane using a semi-dry blot procedure in combination with the Towbin blotting kit (Serva) following the manufacturer’s instructions. Subsequently, membranes were blocked in 1x BlueBlock-reagent (Serva) for 1 h. Incubation with the following primary antibodies occurred over night: rabbit-anti-DNMT1 (#70-201, BioAcademia, 1:1000), mouse-anti-β-ACTIN (#sc-69879, Santa Cruz Biotechnology, 1:1000), rabbit-anti-LC3B (#2775S, Cell Signaling Technology, 1:1000). The following secondary antibodies coupled to horseradish peroxidase (HRP) were applied for 1 h: sheep anti-mouse (#NA9310V, GE Healthcare, 1:4000), donkey anti-rabbit (#NA9340V, GE Healthcare, 1:4000). Membranes were washed after each incubation step three times for 5 min in TBS-T buffer (25 mM Tris/HCl pH 7.5, 137 mM NaCl, 2.7 mM KCl, 0.05% Tween-20). Chemo-luminescent detection of protein bands was performed at the blot documentation system (ChemDoc, BioRad) after applying the HRP substrate solution (Serva). Protein levels were normalized to β-actin.

### Plasmid information

pLAMP1-mCherry [82] was a gift from Amy Palmer (Addgene plasmid #45147; http://n2t.net/addgene:45147; RRID:Addgene_45147). CD63-pEGFP C2 was a gift from Paul Luzio (Addgene plasmid # 62964; http://n2t.net/addgene:62964; RRID:Addgene_62964). 1x GFP-pEGFP N3-HTT was a gift from Mukhran Khundadze (Institute of Human Genetics, University Hospital Jena, Germany).

### Sequencing data analysis

The RNA sequencing data are published in Pensold *et al*. (2020) [12]. Briefly, for isolation of fluorescently labeled cells from adult *Pvalb-Cre/tdTomato/Dnmt1* WT as well as *Pvalb-Cre/tdTomato/Dnmt1* loxP^2^ mice, neurons were from whole brain hemispheres. Following the addition of DAPI, cells were sorted using an ARIA III FACS sorter (BD Biosciences, U.S.A.). RNA was isolated using the TRIzol™ (Invitrogen) protocol according to manufacturer’s instructions. RNA quality was assessed by measuring the RIN (RNA Integrity Number) using the fragment analyzer from Advanced Analytical (U.S.A.). Library preparation for RNA-Seq was performed using the TruSeq™ RNA Sample Prep Kit v2 (Illumina, Cat. N°RS-122-2002, U.S.A.) starting from 50 ng of total RNA. Accurate quantitation of cDNA libraries was performed by using the QuantiFluor™ dsDNA System (Promega, U.S.A.). The size range of final cDNA libraries was determined applying the DNA chip on the fragment analyzer (average 350 bp; Advanced Analytical). cDNA libraries were amplified and sequenced by using the cBot and HiSeq2000 from Illumina (SR; 1×50 bp; ∼30-40 million reads per sample). Sequence images were transformed with Illumina software BaseCaller to bcl files, which were demultiplexed to fastq files with CASAVA v1.8.2. Quality check was done via fastqc (v. 0.10.0, Babraham Bioinformatics, G.B.). Read alignment was performed using STAR (v2.3.0; [83]) to the mm10 reference genome. Data were converted and sorted by samtools 0.1.19 and reads per gene were counted via htseq version 0.5.4.p3. Normalization of raw counts and differential gene expression analysis were performed using the DESeq2 R package (v 1.12.3; [84]). Genes were considered differentially expressed with a Benjamini-Hochberg adjusted P value P < 0.05. Gene lists were submitted to the Database for Annotation, Visualization and Integrated Discovery (DAVID, https://david.ncifcrf.gov) for Gene Ontology enrichment analysis. Results of GO enrichment analysis were visualized in a bar diagram including the respective Benjamini-Hochberg corrected p-value and the number of genes. Heatmaps were generated using R package pheatmap (https://CRAN.R-project.org/package=pheatmap).

### Microscopy and data analysis

For detection of CD63-GFP and LAMP1-mCherry proteins in N2a cells in the endosomal and lysosomal vesicle trafficking assay, cells were imaged using the DMi8 inverted microscope with thunder imaging platform (Leica) with 40x oil objective and incubation (37° C, 5% CO2). Brightfield, FITC- and TRITC-channels were used to check the transfection success of cells before imaging.. Both plasmid conditions were imaged with z-stacks (5 stacks). One complete stack was imaged every 600 ms for a total time frame of 90 s. Endosomal and lysosomal vesicle movement speed, transport direction and distances were tracked and analyzed with the ImageJ software (version 1.52p). CB and N2a cells in the Huntingtin cytotoxicity assay were imaged with the DMi8 inverted microscope with thunder imaging platform (Leica) with 20x objective and incubation (33°C for CB cells, 37 °C for N2a cells, both 5 % CO_2_). Cells were imaged for 6 h or 12 h, capturing an image every 30 min. Huntingtin-GFP was detected in the FITC-channel, and cells were additionally imaged in brightfield. ImageJ (version 1.52p) was used for the analysis of cell survival and tracking of Huntingtin-GFP signal accumulation. Photoshop CC (version 21.2) was applied for image illustration. Graphs and boxplots were generated using Microsoft Excel (version 2019). Significance was analyzed with two-sided *Student’s* t-test or two-way ANOVA. Significance levels: P value < 0.05 *; P value < 0.01 **; P value < 0.001 ***. If not stated differently, experiments were repeated three times.

## Data and materials availability

RNA-sequencing data of FAC-sorted *Pvalb-Cre/tdTomato/Dnmt1* WT and *Pvalb-Cre/tdTomato/Dnmt1 loxP*^2^ samples will be provided on GEO. All other data is available in the main text or the supplementary materials.

## Supplementary Materials

Supplementary materials can be found at www.mdpi.com/xxx/s1.

## Author Contributions

CB designed and performed experiments, data analysis, figure illustration, manuscript writing, manuscript corrections. GP performed experiments, data analysis, figure illustration, manuscript corrections, assisted in manuscript writing. NH performed experiments, data analysis. JL performed experiments, figure illustration, manuscript corrections. JR method establishment, figure illustration, manuscript corrections. DP data analysis and manuscript corrections. GZB conceptual design of the study, manuscript writing.

## Acknowledgments

none.

