## Supplementary figures and images for "The DNA methyltransferase 1 (DNMT1) acts on neurodegeneration by modulating proteostasis-relevant intracellular processes"

### Supplementary Figure S1

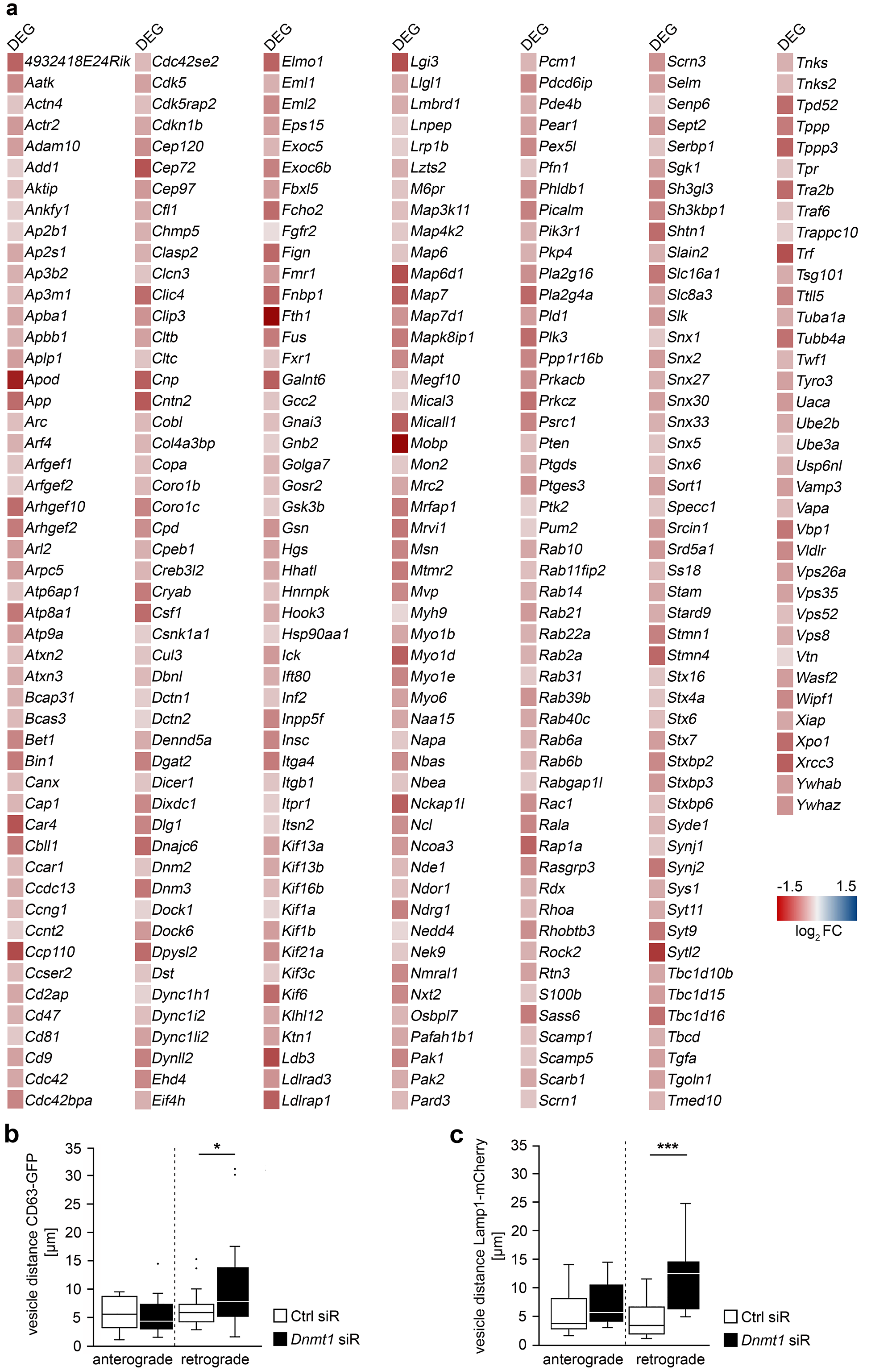

### Supplementary Figure S2

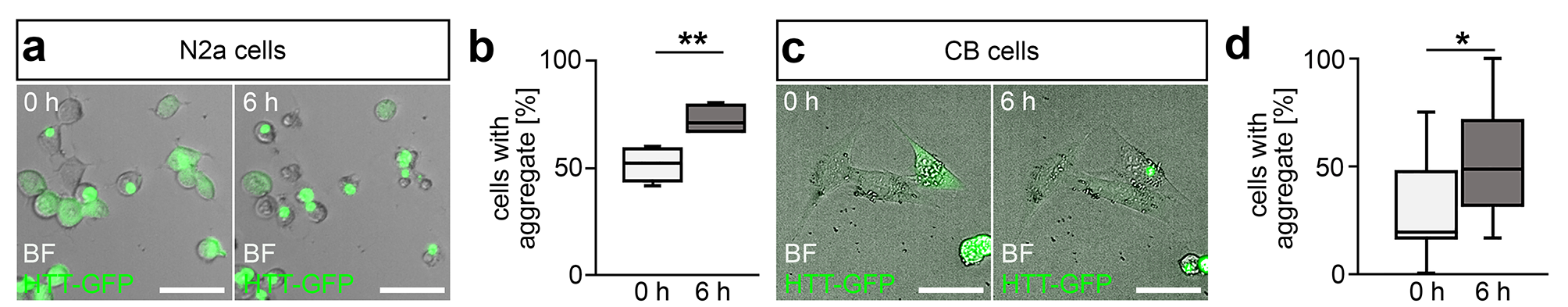
