## Supplementary Legends for "The DNA methyltransferase 1 (DNMT1) acts on neurodegeneration by modulating proteostasis-relevant intracellular processes"

**Figure S1**

(**a**) Heat-map of significantly up-regulated genes in the *Pvalb*-*Cre*/*tdTomato*/*Dnmt1* KO compared to WT FAC-sorted cortical interneurons associated to the GO term *microtubule-based process*, *vesicle-mediated transport* and *perinuclear region of cytoplasm*. Expression level determined by RNA sequencing (*P* < 0.05, Benjamini-adjusted, n = 9 WT and n = 12 KO mice). (**b**) Boxplot showing the distances covered by antero- and retrograde moving CD63-GFP-positive particles of N2a cells treated with either Ctrl siRNA (n = 14 cells, 34 vesicles) or *Dnmt1* siRNA (n= 14 cells, 38 vesicles) and monitored during the endo-lysosomal vesicle tracking. (Two-sided Student’s t-test, * P < 0.05). (**c**) Boxplot showing the distances covered by antero- and retrograde moving LAMP1-mCherry-positive particles of N2a cells treated with either Ctrl siRNA (n = 20 cells, 38 vesicles) or *Dnmt1* siRNA (n= 17 cells, 37 vesicles) and monitored during the endo-lysosomal vesicle tracking. (Two-sided Student’s t-test, *** P < 0.001).

DEG = differentially expressed genes, FC = foldchange, siR = siRNA.

**Figure S2**

(**a**) Representative microphotographs of control N2a cells expressing GFP-labeled mutant HTT (green) showing the increase of cells with aggregates from timepoint 0 h to 6 h. (**b**) Quantification of cell numbers shown in (**a**) (N = 4 experiments, n = 123 cells; Two-sided Student’s t-test, ** P < 0.01). (**c**) Representative microphotographs of control CB cells expressing GFP-labeled mutant HTT (green) showing the increase of cells with aggregates from timepoint 0 h to 6 h. (**d**) Quantification of cell numbers shown in (**c**) (n = 113 cells; Two-sided Student’s t-test, * P < 0.05).

N2a = neuroblastoma cells, CB = cerebellar granule cells, BF = bright field, HTT = Huntingtin. Scale bars: 50 µm in (a, c).
